# ViralMSA: Massively scalable reference-guided multiple sequence alignment of viral genomes

**DOI:** 10.1101/2020.04.20.052068

**Authors:** Niema Moshiri

## Abstract

**Motivation:** In molecular epidemiology, the identification of clusters of transmissions typically requires the alignment of viral genomic sequence data. However, existing methods of multiple sequence alignment scale poorly with respect to the number of sequences.

**Results:** ViralMSA is a user-friendly reference-guided multiple sequence alignment tool that leverages the algorithmic techniques of read mappers to enable the multiple sequence alignment of ultra-large viral genome datasets. It scales linearly with the number of sequences, and it is able to align tens of thousands of full viral genomes in seconds.

**Availability:** ViralMSA is freely available at https://github.com/niemasd/ViralMSA as an open-source software project.

## 1 Introduction

Real-time or near real-time surveillance of the spread of a pathogen can provide actionable information for public health response (Poon *et al*., 2016). Though there is currently no consensus in the world of molecular epidemiology regarding a formal definition of what exactly constitutes a “transmission cluster” (Novitsky *et al*., 2017), all current methods of inferring transmission clusters require a multiple sequence alignment (MSA) of the viral genomes: distance-based methods of transmission clustering require knowledge of homology for accurate distance measurement (Pond *et al*., 2018), and phylogenetic methods of transmission clustering require the MSA as a precursor to phylogenetic inference (Balaban *et al*., 2019; Rose *et al*., 2017; Ragonnet-Cronin *et al*., 2013; Prosperi *et al*., 2011).

The standard tools for performing MSA such as MAFFT (Katoh & Standley, 2013), MUSCLE (Edgar, 2004), and Clustal Omega (Sievers & Higgins, 2014) are prohibitively slow for real-time pathogen surveillance as the number of viral genomes grows. For example, during the COVID-19 pandemic, the number of viral genome assemblies available from around the world grew exponentially in the initial months of the pandemic, but MAFFT, the fastest of the aforementioned MSA tools, scales quadratically with respect to the number of sequences.

In the case of closely-related viral sequences for which a high-confidence reference genome exists, MSA can be accelerated by independently comparing each viral genome in the dataset against the reference genome and then using the reference as an anchor to merge the individual alignments into a single MSA.

Here, we introduce ViralMSA, a user-friendly open-source MSA tool that utilizes read mappers such as Minimap2 (Li, 2018) to enable the reference-guided alignment of ultra-large viral genome datasets.

## 2 Related work

VIRULIGN is another reference-guided MSA tool designed for viral genomes (Libin *et al*., 2019). While VIRULIGN also aims to support MSA of large sequence datasets, its primary objective is to produce codon-correct MSAs (i.e., avoiding frameshifts), whereas ViralMSA’s objective is to produce MSAs for use in real-time applications such as transmission clustering. Thus, while ViralMSA is not guaranteed to yield codon-correct alignments, it is orders of magnitude faster than VIRULIGN, which is critical for rapidly-growing epidemics. Further, because it is codon-correct, VIRULIGN is appropriate for coding regions, whereas ViralMSA is appropriate for whole genomes. Lastly, VIRULIGN requires a thorough annotation of the reference genome, which may be difficult to obtain (especially towards the beginning of a novel outbreak) and does not provide the user to easily utilize different reference genomes for different viral strains. ViralMSA, on the other hand, only requires the reference genome assembly’s GenBank accession number and can build any required index files on-the-fly.

## 3 Results and discussion

ViralMSA is written in Python 3 and is thus cross-platform. ViralMSA depends on BioPython (Cock *et al*., 2009) and whichever read mapper the user chooses, which is Minimap2 by default (Li, 2018). In addition to Minimap2, ViralMSA supports STAR (Dobin *et al*., 2013), Bowtie 2 (Langmead & Salzberg, 2012), and HISAT2 (Kim *et al*., 2019), though the default of Minimap2 is strongly recommended: Minimap2 is much faster than the others (Li, 2018) and is the only mapper that consistently succeeds to align all genome assemblies against an appropriate reference across multiple viruses. ViralMSA’s support for read mappers other than Minimap2 is primarily to demonstrate that ViralMSA is flexible, meaning it will be simple to incorporate new read mappers in the future.

ViralMSA takes the following as input: (1) a FASTA file containing the viral genomes to align, (2) the GenBank accession number of the reference genome, and (3) the mapper to utilize (Minimap2 by default). ViralMSA will pull the reference genome from GenBank and generate an index using the selected mapper, both of which will be cached for future alignments of the same viral strain, and will then execute the mapping. ViralMSA will then process the results and output an MSA in the FASTA format. For commonly-studied viruses, the user can simply provide the name of the virus instead of an accession number, and ViralMSA will select an appropriate reference genome. The user can also choose to provide a local FASTA file containing a reference genome, which may be useful if the desired reference does not exist on GenBank or if the user wishes to conduct the analysis offline.

Because it uses the positions of the reference genome as anchors with which to merge the individual pairwise alignments, ViralMSA only keeps matches, mismatches, and deletions with respect to the reference genome: it discards all insertions with respect to the reference genome. For closely-related viral strains, insertions with respect to the reference genome are typically unique and thus lack usable phylogenetic or transmission clustering information, so their removal results in little to no impact on downstream analyses (Tab. 1).

**Table 1.** Multiple sequence alignment accuracy

| Virus | MAFFT (S) | ViralMSA (S) | MAFFT (P) | ViralMSA (P) |
| --- | --- | --- | --- | --- |
| Ebola | 0.9957 | 0.9873 | 0.9998 | 0.9816 |
| HCV | 0.9995 | 0.9506 | 0.9999 | 0.9678 |
| HIV-1 | 0.9786 | 0.9705 | 0.9957 | 0.9941 |

Correlation coefficients are shown for Mantel tests between curated “ground truth” MSAs and those estimated by MAFFT and ViralMSA. *S* and *P* denote Spearman and Pearson Correlation, respectively. 1 indicates perfect correlation, -1 indicates perfect anticorrelation, and 0 indicates no correlation.

In order to test MSA accuracy, we obtained curated Ebola, HCV, and HIV-1 full genome MSAs from the Los Alamos National Laboratory (LANL) Sequence Databases, which we used as our ground truth. In order to benchmark MSA runtime, we obtained a large collection of SARS-CoV-2 complete genomes from the Global Initiative on Sharing All Influenza Data (GISAID) database. We then used both MAFFT and ViralMSA to estimate MSAs. No other MSA tools were included in the comparison due to being orders of magnitude slower than MAFFT (which would render the full experiment infeasible).

Because ViralMSA’s objective is to be utilized in real-time applications such as transmission clustering workflows, which typically rely on pairwise distances between samples, we computed pairwise sequence distances between every pair of samples in each MSA under the TN93 model of sequence evolution (Tamura & Nei, 1993) using the pairwise distance calculator implemented in HIV-TRACE (Pond et al., 2018). Then, for each estimated MSA, we measured alignment accuracy by computing the Mantel correlation test between the curated (“true”) and estimated MSAs. In order to measure performance, we subsampled the full SARS-CoV-2 dataset, with 10 replicates for each dataset size. We then computed MSAs of each subsampled alignment replicate.

As can be seen, ViralMSA in its default mode is consistently orders of magnitude faster than MAFFT (Figs. 1, S1), yet it produces MSAs with just slightly lower accuracy, even across different viruses (Tab. 1) and different levels of subsampling (Figs. S2-S3). Further, both MAFFT and ViralMSA produce MSAs that tend to underestimate TN93 distance, with ViralMSA underestimating slightly more significantly (Fig. S4). Note that ViralMSA’s speed and accuracy stem from the algorithmic innovations of the selected read mapper (not from ViralMSA itself), meaning ViralMSA can natively improve as read mapping tools evolve.

**Fig. 1.**
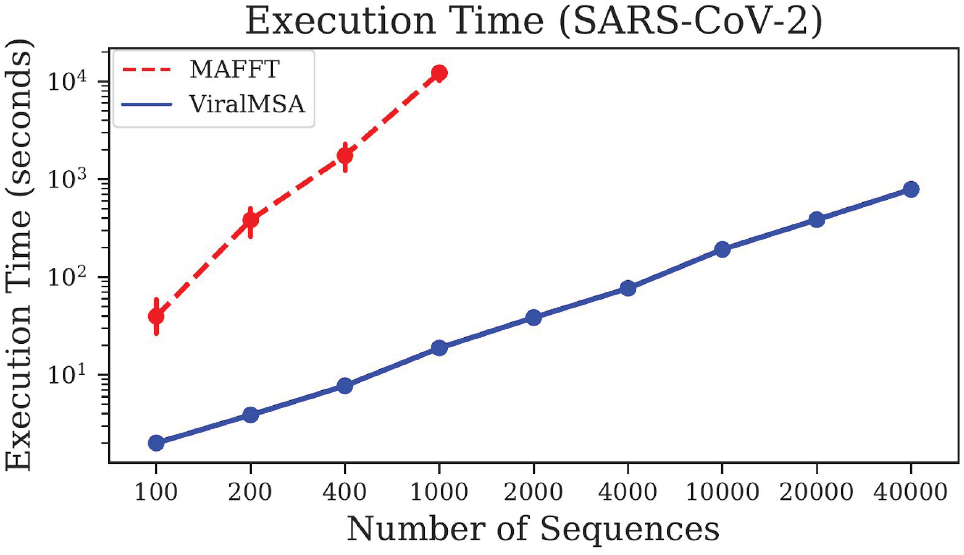
Execution time. Execution time (seconds) for SARS-CoV-2 MSAs (genome length 29kb) estimated by MAFFT and ViralMSA for various dataset sizes. All runs were executed sequentially on an 8-core 2.0 GHz Intel Xeon CPU with 30 GB of memory.

## Acknowledgements

We would like to thank Heng Li and his exceptional work in developing Minimap2. His work is integral to ViralMSA’s speed and accuracy.

## References

Balaban, M. et al.(2019) TreeCluster: Clustering biological sequences using phylogenetic trees. PLoS One, 14(8), e0221068.

Cock, P.J.A. et al. (2009) Biopython: freely available Python tools for computational molecular biology and bioinformatics. Bioinformatics, 25(11), 1422–1423.

Dobin, A. et al. (2009) STAR: ultrafast universal RNA-seq aligner. Bioinformatics, 29(1), 15–21.

Edgar, R.C. (2004) MUSCLE: a multiple sequence alignment method with reduced time and space complexity. BMC Bioinform., 5, 113.

Langmead, B. and Salzberg, S.L. (2012) Fast gapped-read alignment with Bowtie 2. Nat. Methods, 9, 357–359.

Kim, D. et al. (2019) Graph-based genome alignment and genotyping with HISAT2 and HISAT-genotype. Nat. Biotechnol., 37, 907–915.

Li, H. (2018) Minimap2: pairwise alignment for nucleotide sequences. Bioinformatics, 34(18), 3094–3100.

Libin, P.J.K. (2019) VIRULIGN: fast codon-correct alignment and annotation of viral genomes. Bioinformatics, 35(10), 1763–1765.

Katoh, K. and Standley, D.M. (2013) MAFFT Multiple Sequence Alignment Software Version 7: Improvements in Performance and Usability. Mol. Biol. Evol., 30(4), 772–780.

Novitsky, V. et al. (2017) Phylogenetic Inference of HIV Transmission Clusters. Infect. Dis. Transl. Med., 3(2), 51–59.

Pond, S.L.K. et al. (2018) HIV-TRACE (TRAnsmission Cluster Engine): a Tool for Large Scale Molecular Epidemiology of HIV-1 and Other Rapidly Evolving Pathogens. Mol. Biol. Evol., 35(7), 1812–1819.

Poon, A.F.Y. et al. (2016) Near real-time monitoring of HIV transmission hotspots from routine HIV genotyping: an implementation case study. Lancet HIV, 3(5), e231–e238.

Prosperi, M.C.F. et al. (2011) A novel methodology for large-scale phylogeny partition. Nat. Commun., 2, 321.

Ragonnet-Cronin, M. et al. (2013) Automated analysis of phylogenetic clusters. BMC Bioinform., 14, 317.

Rose, R. et al. (2017) Identifying Transmission Clusters with Cluster Picker and HIV-TRACE. AIDS Res. Hum. Retroviruses, 33(3), 211–218.

Sievers, F. and Higgins, D.G. (2014) Clustal Omega, accurate alignment of very large numbers of sequences. Methods Mol. Biol., 1079, 105–116.

Tamura, K. and Nei, M. (1993) Estimation of the number of nucleotide substitutions in the control region of mitochondrial DNA in humans and chimpanzees. Mol. Biol. Evol., 10(3), 512–526.

