## Supplementary Materials for "ViralMSA: Massively scalable reference-guided multiple sequence alignment of viral genomes"

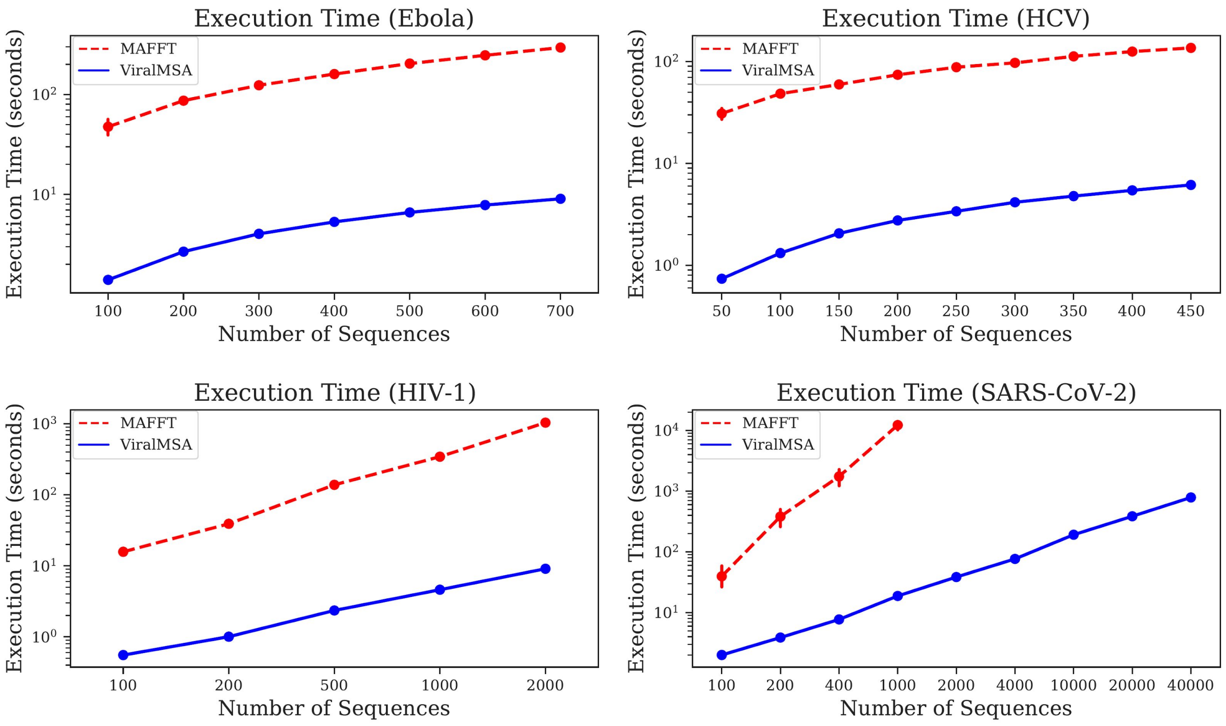


**Fig. S1. Execution time.** Execution time (seconds) is shown for Ebola, HCV, HIV-1, and SARS-CoV-2 MSAs estimated by MAFFT and ViralMSA for various dataset sizes.

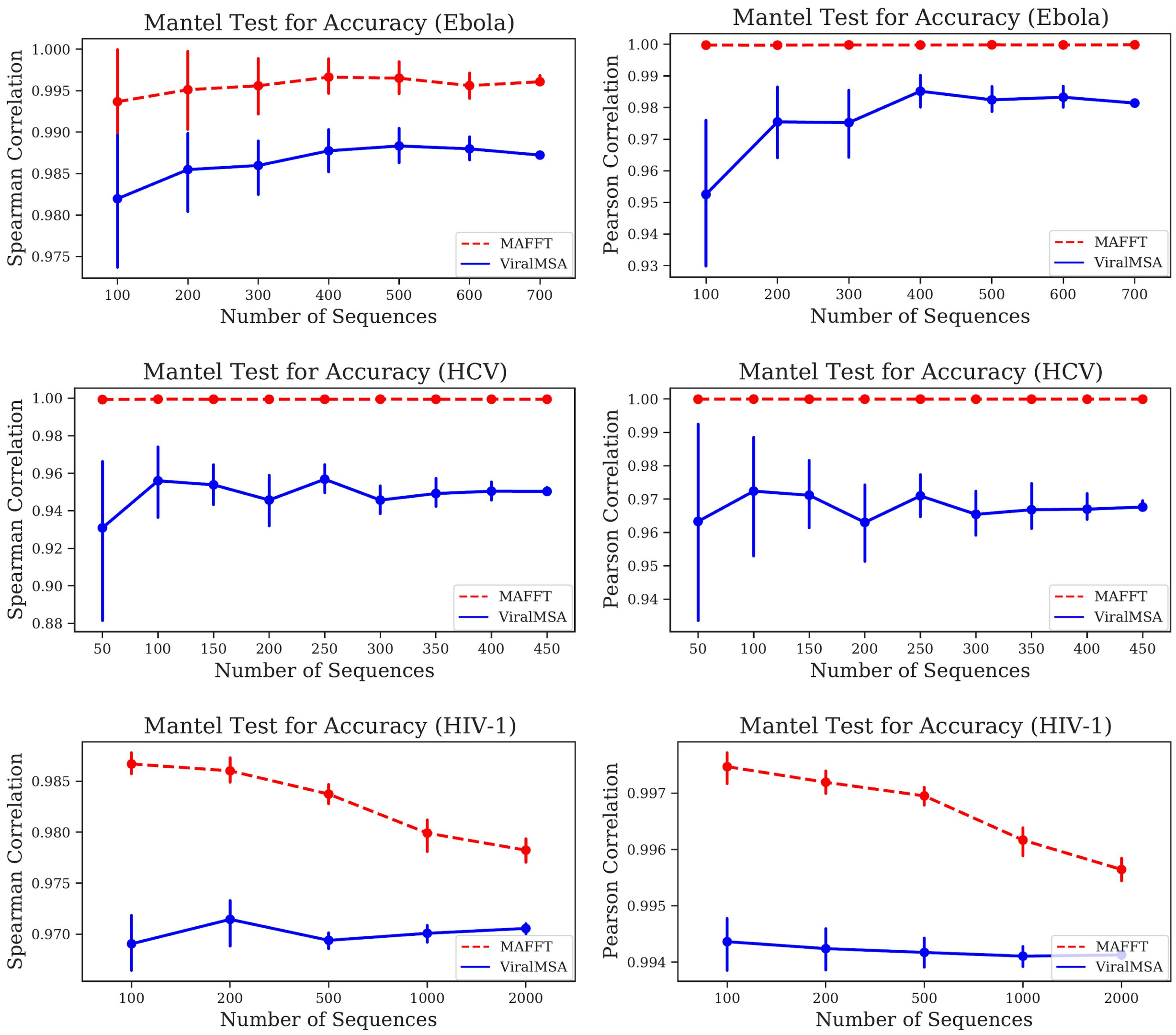


**Fig. S2. MSA accuracy.** Spearman and Pearson correlation coefficients are shown for Mantel tests between curated "ground truth" MSAs and those estimated by MAFFT and ViralMSA for Ebola, HCV, and HIV-1 datasets of various sizes. 1 indicates perfect correlation, -1 indicates perfect anticorrelation, and 0 indicates no correlation.

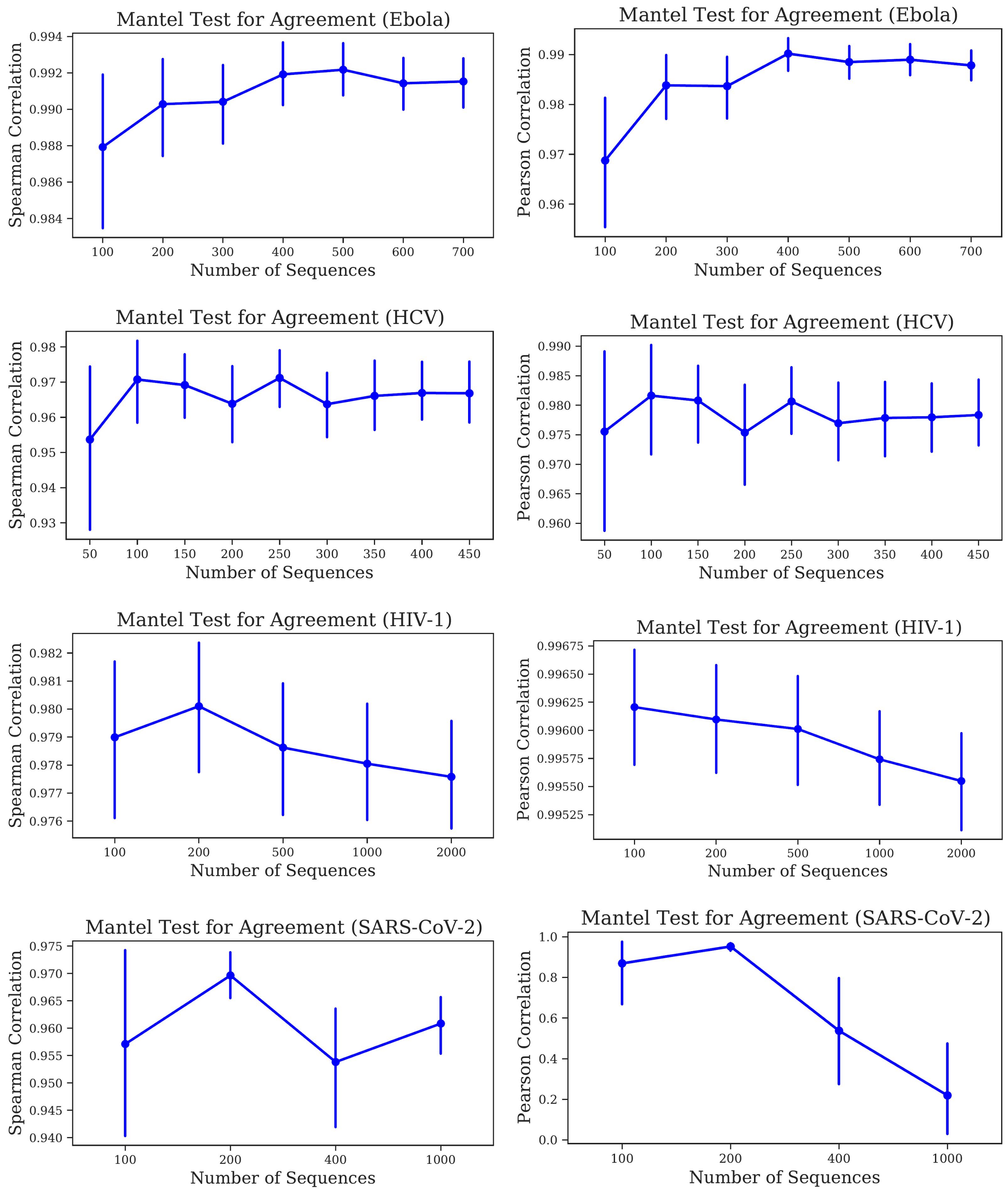


**Fig. S3. MSA agreement.** Spearman and Pearson correlation coefficients are shown for Mantel tests between MSAs estimated by MAFFT and those estimated by ViralMSA for Ebola, HCV, HIV-1, and SARS-CoV-2 datasets of various sizes. 1 indicates perfect correlation, -1 indicates perfect anticorrelation, and 0 indicates no correlation.

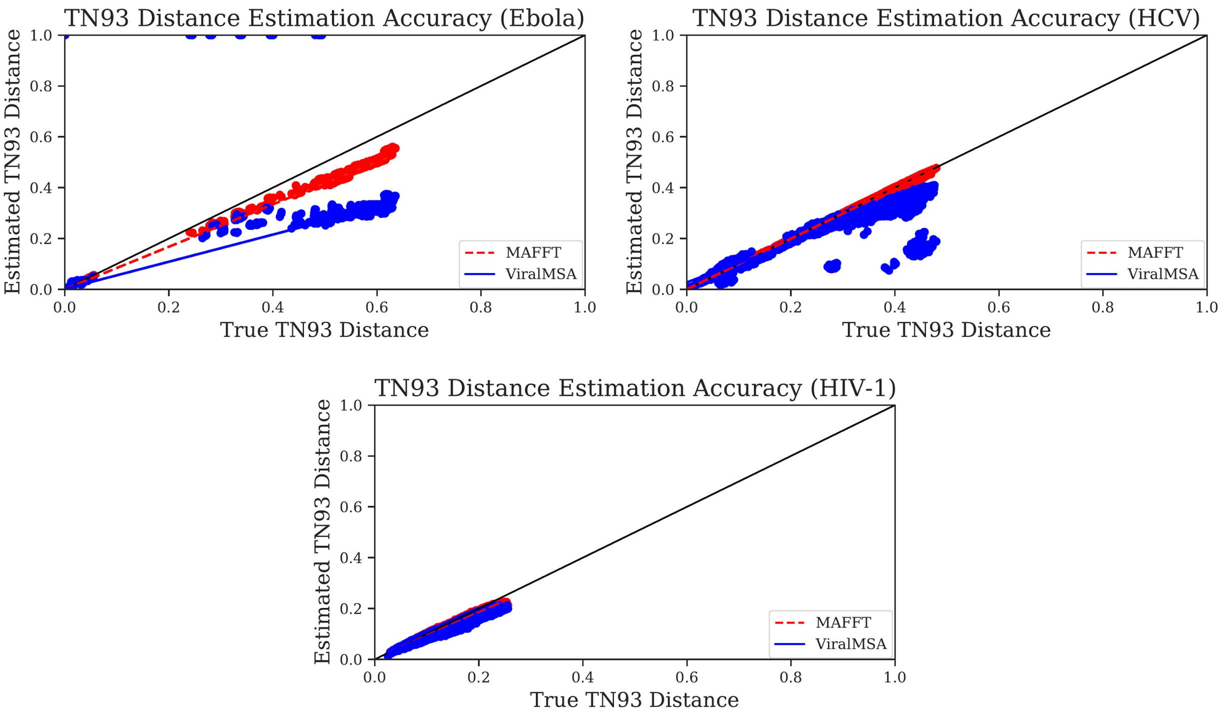


**Fig. S4. Distance estimation accuracy.** TN93 distances computed from MSAs estimated by MAFFT and ViralMSA ($y$-axis) are shown against the corresponding "true" TN93 distances computed from curated MSAs ($x$-axis) for Ebola, HCV, and HIV-1. The line $y = x$ (denoting perfect estimation) is shown in black.
